# Comparative genomics reveals the diversity of CRISPR-Cas systems among neonatal sepsis causing group B *Streptococcus agalactiae*

**DOI:** 10.1101/2023.09.27.559688

**Authors:** Sudeep D Ghate, R. Shyama Prasad Rao, Rajesh P Shastry, Deepak Pinto, Praveenkumar Shetty

## Abstract

The pathogen *Streptococcus agalactiae*, or Group B *Streptococcus* (GBS) infection is the leading cause of neonatal sepsis and meningitis in neonates. In this study, we aimed to investigate the occurrence and diversity of the CRISPR-Cas system in *S. agalactiae* genomes using computational biology approaches. A total of 51 out of 52 complete genomes (98.07%) of *S. agalactiae* possess CRISPR arrays (75 CRISPR arrays) with 17 strains possessing multiple CRISPR arrays. There were only two CRISPR-Cas systems – type II-A system and type I-C system in *S. agalactiae* strains. RNA secondary structure analysis through direct repeat analysis showed that the analyzed strains could form stable secondary structures. The 16S rRNA phylogeny exhibited clustering of the strains into three major clades grouped on the type of CRISPR-Cas system. The anti-CRISPRs that contribute to CRISPR-Cas system diversity and prevent genome editing were also detected. These results provide valuable insights into elucidating the evolution, diversity, and function of CRISPR/Cas elements in this pathogen.

## INTRODUCTION

*Streptococcus agalactiae* (also known as Group B *Streptococcus*, GBS), a Gram-positive bacterium, is a common commensal of intestinal and reproductive tracts in healthy adults. It can be transmitted from mother to newborn during birth [1]. The GBS is a cause of stillbirth, chorioamnionitis, and neonatal infections including pneumonia, bacteremia, and meningitis. GBS-sepsis has a mortality rate of 10-18%, with a colonization rate of approximately 18–35% in pregnant women, and neonatal infection rates of 0.4 to 1.1 cases per 1000 live births [2,3]. The infection process is mediated by multifunctional GBS virulence factors that could be a challenge to the immune-deficient neonates [4]. GBS displays virulence factors including a potent hemolytic toxin, proteases, and multiple surface proteins to conquer host tissues [5].

Prokaryotes employ CRISPR-Cas systems (clustered regularly interspaced short palindromic repeat, with CRISPR-associated proteins), which provide sequence-based adaptive immunity against invasive transposable elements, conjugative plasmids, and phages [6]. About 40% of bacteria and 90% of archaea are equipped with CRISPR-Cas systems. Moreover, microbes may have more than one type of CRISPR–Cas system which function towards specific template based recognition, targeting, and degradation of exogenous nucleic acids [7]. These systems could differ in type of Cas proteins present and spacer sequences and also the length and number of CRISPR repeats. Although, initially known for its involvement in viral defense, recent findings suggest involvement of CRISPR-Cas systems in regulation of expression of virulence genes and escape host immunity [8]. CRISPR-Cas systems were earlier considered as adaptive immune system and widely studied in *Streptococcus thermophilus* [9].

Studies suggest that mainly three types of CRISPR-Cas9 systems are employed by Streptococcus sp. type I, type II, and type III. In addition to these, they also harbor a single type V and unknown CRISPR loci.[10]. The recent reports on the emergence of hypervirulent *S. agalactiae* suggest the contribution of phages and other mobile genetic elements (MGE) in adaptation to different hosts and its virulence profile [11]. These phage-associated genes may play a major role in biological success of the strains by acting as delivery vehicles of resistance and virulence genes [12]. Recently, CRISPR analysis has been used as tool to follow maternal GBS colonization and also as a typing technique over traditional subtyping systems [13]. While extensive details are available on *S. thermophilus* and other animal pathogenic streptococci, detailed information on the CRISIPR-Cas systems in human pathogenic *S. agalactiae* are lacking. Therefore, in this regard, we sought to investigate the occurrence and diversity of CRISPR-Cas systems in *S. agalactiae* genomes. We used CRISPRminer2 server[14] and CRISPRCasFinder [15] - two most inclusive and widely used resources for the identification of CRISPR arrays and cas genes. We report here the diversity and provide insights into existing CRISPR-Cas systems in *S. agalactiae* based on 52 complete genomes of GBS of human origin.

## METHODOLOGY

### Sequence selection and retrieval

The data set included complete genomes of *Streptococcus agalactiae*. Only the complete genomes with human/*homo sapiens* as hosts were selected and retrieved from NCBI website (https://www.ncbi.nlm.nih.gov/, last accessed on 21-08-2022). Except for the reference strain NGBS128 none of the other genomes had any plasmid sequences. A total of 52 such sequences were selected and their fasta files downloaded for NCBI.

### Detection of CRISPR-Cas features

The complete genomes of the 52 strains were screened for the presence of complete CRISPR– Cas loci using CRISPRminer2 server. CRISPRminer2 is a comprehensive tool that uses a comparative genomics approach to identify and annotate CRISPR–Cas loci. This tool also helps with multiple detection options, including anti-CRISPR detection and annotation, self-targeting spacer search, repeat type identification, bacteria–phage interaction detection, and prophage detection. Only “confirmed CRISPRs” identified by the CRISPRminer2 tool were selected for further analysis. The strains which then did not show CRISPR loci were eliminated and rest of the strains were retained for further analysis. The results were also corroborated by checking with CRISPRCasFinder.

### Signature genes

The data from CRISPRminer2 was tabulated and a list of signature genes were determined. A tile map was generated to visualise the presence and distribution of these genes amongst the 43 strains. CRISPR map server was also used to obtain in depth information on each of the strains. The CRISPR repeats were analysed through multiple sequence alignment and the aligned direct repeats visualised using the WebLogo program [16].

### RNAFold Webserver

Direct repeats (DR) obtained via CRISPRminer2 were then compiled and 11 unique repeats found were then used to generate free energy structures via the RNAFold server[17]. The RNAfold Webserver set to default parameters was used to predict the RNA secondary structure and minimum free energy (MFE) of each DR.

### Phylogenetic Trees

To understand CRISPR-Cas distribution in the genomes from a phylogenetic perspective, complete 16S rDNA sequences from 52 genomes were retrieved from NCBI and aligned using MUSCLE in MEGAX [18]. ML statistical method with model selection was used to compute BIC score and AICc value of 24 different nucleotide substitution models. A maximum likelihood phylogenetic tree was constructed (Kimura-2 model of nucleotide substitution) and bootstrap analysis with 1000 random replicates. The cas1 and cas9 genes were aligned and a ML phylogenetic tree was constructed with 1000 bootstrap values. *Streptococcus pyogenes* was taken as the outgroup.

### Spacer analysis

The spacer targets were identified using the CRIPSRminer2. The visual representation of the CRISPR spacers was performed using Excel macros, with each unique colour combination representing one unique spacer sequence

## RESULTS

### Sequence selection and retrieval

A search for *S. agalactiae* genomes in the NCBI database listed 1515 sequenced genomes amongst which 128 were complete genomes. Out of these only the ones affecting human/ *homo sapiens* hosts which resulted in a total of 52 genomes were considered for further analysis. Out of these 52 strains, 51 were determined to possess CRISPR arrays with a total of 75 CRISPR arrays detected and 21 strains possessing multiple of these CRISPR arrays. BJ01 and Sag27 were noted to have the highest number of individual arrays each with three CRISPR-Cas arrays.

### Genomic context analysis of confirmed CRISPR-Cas loci

The selected strains were uploaded to CRISPRminer2 web server and results are tabulated in the Table 1. CRISPRminer2 provided details on the number and type of CRISPR locus found, number of spacers and direct repeats (DRs), including DR types, signature genes found within each locus. Other information such as number of prophages, anti-CRISPRs, mobile genetic elements and self-targeting spacers were all obtained from this server. All the strains were classified either into Type II, Type I or orphan CRISPR types (Supplementary table 1). Two types of DRs were found, II having 47 repeats and I having only 12, whilst the remaining 16 repeats were determined to be NA (Not applicable). GBS28 and NGB061 had the highest number of DRs with 31 whilst possessing only 1 CRISPR array. Meanwhile, Sag158 and BJ01 have 31 and 35 DRs respectively but with multiple CRISPR arrays. The individual CRISPR length was observed to have a wide range with FDAARGO_670 and B509 having 7693bp and 7363bp being on the higher end. Meanwhile, BGS-M002 has the shortest CRISPR with only 102 bp. Two types of Anti CRISPR (Acr) regions were also detected with AcrIIA21 being 108 aa long and being present in 26 strains whilst AcrIIA18 being 176 aa in length and present in just GBS1-NY. NGBS061 and BJ01 both possess the highest amount of self-targeting spacers with 11 each. Strain NGBS128 was found to have the greatest number of Mobile Genetic Elements (MGEs) with 37 in its single CRISPR array.

**Table 1:** Genomic properties and CRISPR-Cas type of the 52 GBS strains used in this study.

| # | Strain/Accession number | Source | Condition | CRISPR type | Length (Mb) | GC% | Protein Count |
| --- | --- | --- | --- | --- | --- | --- | --- |
| 1 | 32790-3A / NZ_CP029561.1 | Guangzhou, China | Hospital | II-A | 2.15 | 35.7 | 2167 |
| 2 | 874391 / NZ_CP022537.1 | Japan | Vagina | II-A | 2.15 | 35.5 | 1991 |
| 3 | B111 / NZ_CP021772.1 | Shenzhen, China | Neonatal sepsis | II-A | 2.15 | 35.4 | 2021 |
| 4 | B507 / NZ_CP021771.1 | Shenzhen, China | Vagina (mother) | II-A, I-C | 2.08 | 35.4 | 1936 |
| 5 | B508 / NZ_CP021770.1 | Shenzhen, China | Vagina (mother) | II-A | 2.20 | 35.6 | 2204 |
| 6 | BJ01 / NZ_CP059383.1 | Beijing, China | Neonate blood | Orphan | 2.15 | 35.7 | 2037 |
| 7 | CJB111 / NZ_CP063198.2 | USA | Blood | II-A | 2.09 | 35.5 | 1955 |
| 8 | CNCTC 10_84/ NZ_CP006910.1 | Atlanta, USA | Hospital | II-A | 2.01 | 35.4 | 2046 |
| 9 | COH1 / NZ_HG939456.1 | Institute Pasteur | Sepsis (new-born) | II-A | 2.07 | 35.4 | 1893 |
| 10 | CU_GBS_08 / NZ_CP010874.1 | Hong Kong | Hospital | II-A, I-C | 2.08 | 35.4 | 1987 |
| 11 | CU_GBS_98 / NZ_CP010875.1 | Hong Kong | Meningitis (Hospital) | II-A, I-C | 2.03 | 35.4 | 1916 |
| 12 | CUGBS591 / NZ_CP021862.1 | Hong Kong | Arthritis (Hospital) | II-A, I-C | 2.23 | 35.8 | 2103 |
| 13 | GBS11 / NZ_CP041999.1 | Houston, USA | Blood | II-A | 2.14 | 35.6 | 2180 |
| 14 | GBS19 / NZ_CP042000.1 | Houston, USA | Blood | II-A | 2.10 | 35.5 | 2120 |
| 15 | GBS1-NY / NZ_CP007570.1 | USA | Blood | II-A | 2.24 | 35.9 | 2059 |
| 16 | GBS28 / NZ_CP042001.1 | Tennessee, USA | Health Centre | II-A | 2.14 | 35.7 | 2024 |
| 17 | GBS2-NM / NZ_CP007571.1 | USA | Hospital | II-A | 2.21 | 35.9 | 2036 |
| 18 | GBS30 / NZ_CP042002.1 | Houston, USA | Blood | II-A | 2.08 | 35.5 | 2096 |
| 19 | GBS6 / NZ_CP007572.1 | Houston, USA | Hospital | II-A | 2.23 | 35.8 | 2054 |
| 20 | GBS7 / NZ_CP041998.1 | Houston, USA | Blood | II-A | 2.09 | 35.5 | 1957 |
| 21 | GBS85147 / NZ_CP010319.1 | Rio de Janeiro, Brazil | New-born | II-A | 2.00 | 35.5 | 1992 |
| 22 | GBS-M002 / NZ_CP013908.1 | Taiwan | Cervix | II-A | 2.09 | 35.6 | 1955 |
| 23 | H002 / NZ_CP011329.1 | Guangxi, China | Vagina | II-A | 2.15 | 35.7 | 1984 |
| 24 | HU-GS5823 / NZ_AP018935.1 | Hokkaido, Japan | Hospital | II-A | 2.23 | 35.6 | 2233 |
| 25 | NEM316 / NC_004368.1 | Institute Pasteur | Septicemia | II-A | 2.21 | 35.6 | 2227 |
| 26 | NGBS061 / NZ_CP007631.2 | Toronto, Canada | Health Centre | II-A | 2.22 | 35.5 | 2275 |
| 27 | NGBS572 / NZ_CP007632.1 | Toronto, Canada | Health Centre | II-A, I-C | 2.06 | 35.5 | 2079 |
| 28 | PLGBS13 /<br>NZ_CP029749.1 | Alberta, Canada | Wound (Soft<br>tissue) | II-A, I-C | 2.10 | 35.5 | 2122 |
| 29 | Sag153 /<br>NZ_CP036376.1 | Shanghai, China | Vagina | II-A, I-C | 2.17 | 35.8 | 2223 |
| 30 | Sag158 /<br>NZ_CP019979.1 | Shanghai, China | Hospital | II-A, I-C | 2.10 | 35.7 | 1941 |
| 31 | Sag27 /<br>NZ_CP031556.1 | Shanghai, China | Perianal region | II-A,<br>Orphan<br>(2) | 2.21 | 35.7 | 2074 |
| 32 | Sag37 /<br>NZ_CP019978.1 | Shanghai, China | Blood | II-A, I-C | 2.20 | 35.8 | 2250 |
| 33 | SG-M1 /<br>NZ_CP012419.2 | Singapore | Blood | II-A, I-C | 2.12 | 35.5 | 2180 |
| 34 | SG-M158 /<br>NZ_CP021864.1 | Singapore | Blood | II-A, I-C | 2.11 | 35.5 | 2025 |
| 35 | SG-M163 /<br>NZ_CP021863.1 | Singapore | Blood | II-A, I-C | 2.12 | 35.5 | 2025 |
| 36 | SG-M25<br>NZ_CP021867.1 | Singapore | Blood | II-A,<br>Orphan | 2.21 | 35.7 | 2075 |
| 37 | SG-M29 /<br>NZ_CP021866.1 | Singapore | Blood | II-A, I-C | 2.12 | 35.5 | 2025 |
| 38 | SG-M4 /<br>NZ_CP021870.1 | Singapore | Blood | II-A | 2.07 | 35.5 | 2085 |
| 39 | SG-M50 /<br>NZ_CP021865.1 | Singapore | Blood | II-A, I-C | 2.12 | 35.5 | 2023 |
| 40 | SG-M6 /<br>NZ_CP021869.1 | Singapore | Blood | II-A, I-C | 2.11 | 35.6 | 1954 |
| 41 | SG-M8 /<br>NZ_CP021868.1 | Singapore | Blood | II-A | 2.17 | 35.6 | 2186 |
| 42 | SS1 /<br>NZ_CP010867.1 | Houston, USA | Blood | II-A | 2.09 | 35.5 | 2110 |
| 43 | SS1168 /<br>NZ_CP038809.1 | Houston, USA | Hospital | II-A | 2.04 | 35.4 | 1911 |
| 44 | 2012-845 /<br>CP051842.1 | Versailles, France | Blood | 0<br>CRISPR | 1.53 | 35.3 |  |
| 45 | B105 /<br>NZ_CP021773.1 | Shenzhen, China | Blood sample<br>from a new-born | Orphan | 2.27 | 35.7 | 2076 |
| 46 | B509 /<br>NZ_CP021769.1 | Shenzhen, China | Vagina swab from<br>a perinatal mother | II-A | 2.06 | 35.5 | 1928 |
| 47 | S9968 /<br>NZ_CP058666.1 | Seoul, South Korea | Urine | II-A,<br>Orphan<br>(2) | 2.20 | 35.7 | 2088 |
| 48 | NGBS128 /<br>NZ_CP012480.1 | Greater Toronto<br>area/Peel, Canada | Infection sample | Orphan | 2.08 | 35.7 | 1879 |
| 49 | FDAARGOS_254 /<br>NZ_CP020449.2 | DC, USA | Blood | II-A | 2.22 | 35.7 | 2060 |
| 50 | FDAARGOS_512 /<br>NZ_CP033822.1 | DC, USA | Endotracheal<br>aspirate | Orphan | 2.13 | 35.6 | 2000 |
| 51 | FDAARGOS_669 /<br>NZ_CP044091.1 | DC, USA | Clinical isolate | II-A | 2.07 | 35.4 | 1937 |
| 52 | FDAARGOS_670 /<br>NZ_CP044090.1 | DC, USA | Clinical isolate | II-A | 2.21 | 35.8 | 2098 |

### Cas genes

The tile map generated using the presence/absence matrix shows the distribution of signature genes amongst the 51 genomes (Figure 1). From the given cas genes only cas1, cas2, csn2 (Casein Beta which is a Protein Coding gene) and cas9 were seen to be present in all the 51 strains. The strains B507, CUGBS591, CU_GBS_08, CU_GBS_98, NGBS572, Sag153, Sag37, SG-M1, SG-M158, SG-M63, SG-M29 and SG-M50 possessed 2 CRISPR arrays and hence have the maximum cas proteins. None of the genomes possessed any transposons in the CRISPR loci.

**Figure 1:**
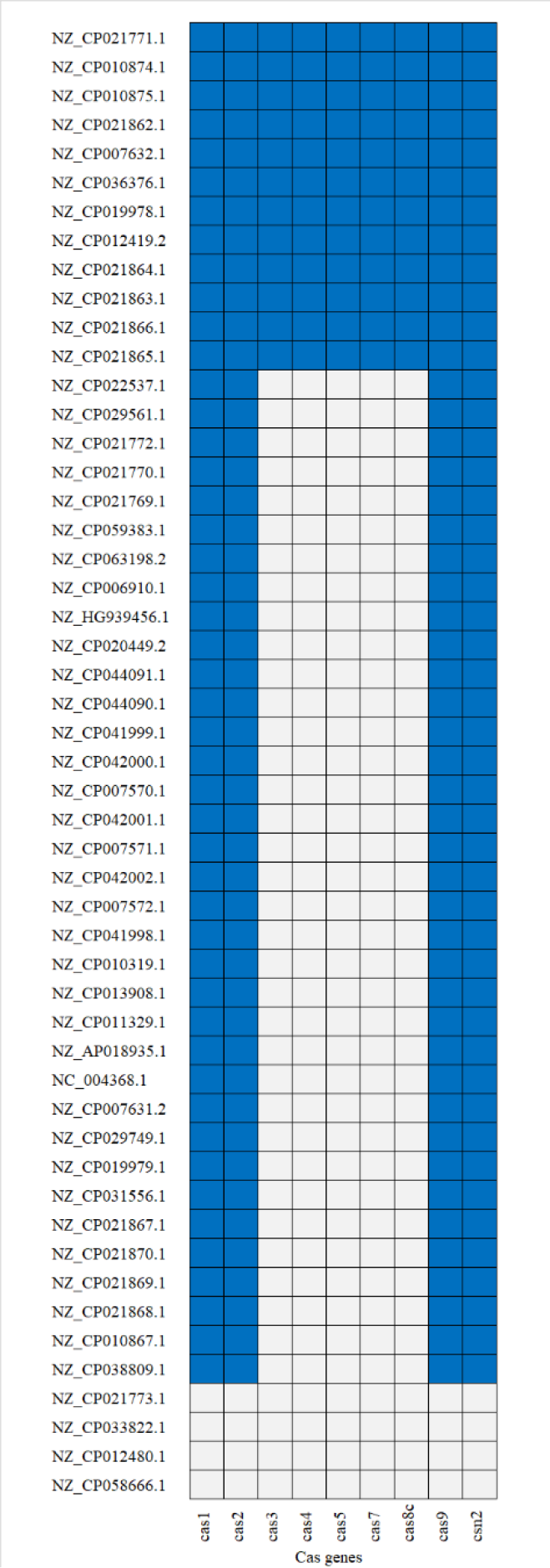
Heatmap of presence/absence of various signature cas genes amongst the 52 strains. The tiles in dark blue denote the presence whilst the ones in light blue show absence.

### Direct repeats

The DRs from all the CRISPRs were collected and the duplicates were removed. The 11 unique repeats were then uploaded to the RNAFold Webserver from which the free energy structure was obtained as seen in Figure 2. Shorter or incomplete DR sequences were eliminated and the remaining 9 structures were taken into consideration. DR1 and DR2 are seen to have the highest minimum free energy (MFE) value whilst DR2 has the lowest making it the most stable out of the 9 DR structures. The MFE of ribonucleic acids (RNAs) increases at an apparent linear rate with sequence length and the lower the MFE, the more stable the structure [19]. In this case DR2 with -13.10 kcal/mol is seen to be the most stable out of the 9 predicted structures. Both DR1 and DR3 have the least stable structure with -0.3 kcal/mol and -0.4 kcal/mol respectively.

**Figure 2:**
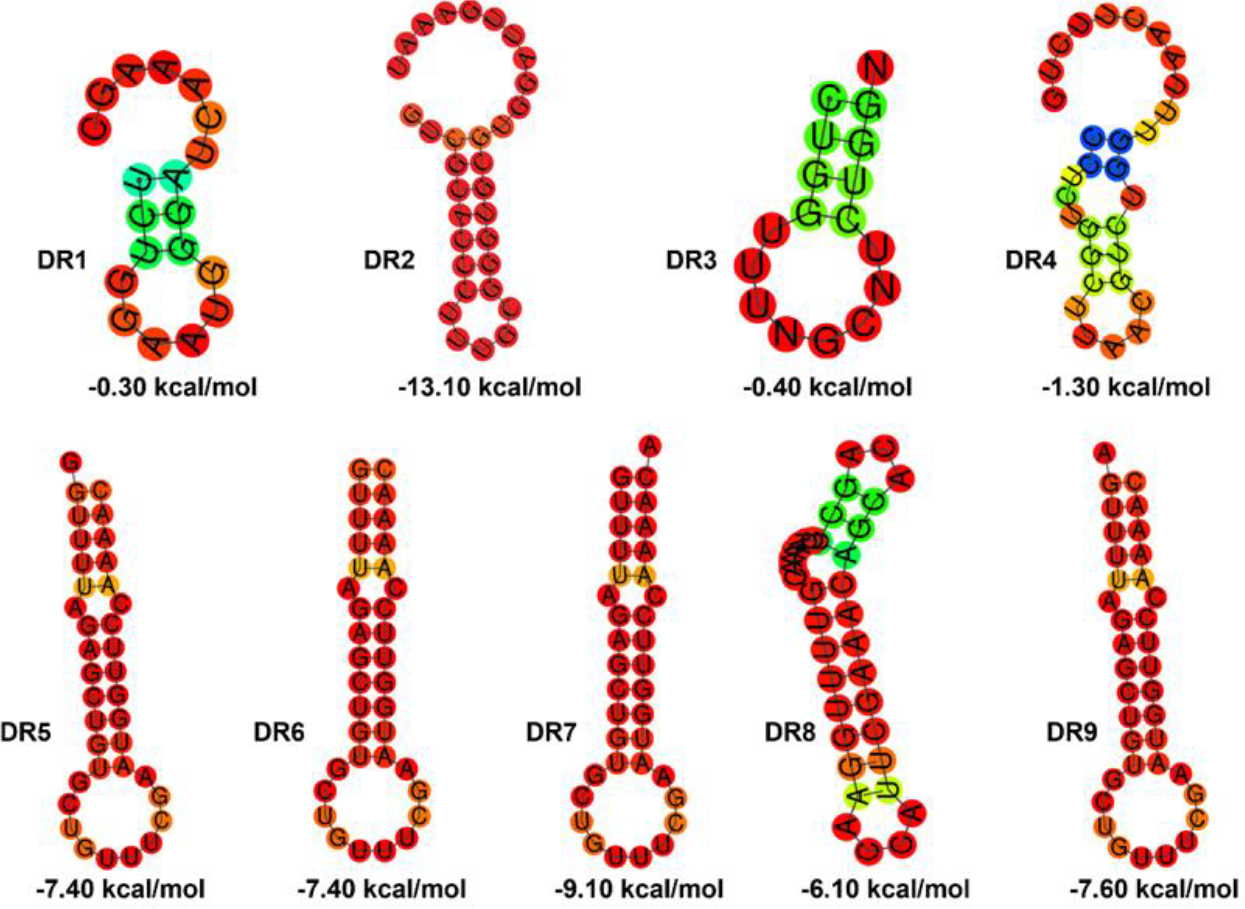
The secondary structure for consensus 11 unique direct repeat sequences of CRISPR arrays in GBS strains. The kcal/mol indicates the minimum free energy (MFE) which is known to increase at a clear linear rate with sequence length. The colours represent the base-pair probability range.

### Spacer analysis

In total, 862 spacers were identified among GBS genomes positive for CRISPR loci. Of the identified spacers, 812 were unique (Supplementary figure 1). The spacers in each array ranged from 23 to 104. Among the genomes, the least number of spacers (1) was seen within the CRISPR locus of BJ01, while the highest number of spacers (31) was seen in GBS28 genome with an average of 11.4 spacers per array. An analysis of spacer sequences showed 212 spacers to match plasmids (24.79%) and 568 spacers (66.43%) to match phages. The CRISPRminer2 prediction indicated the absence of self-targeting spacers. Furthermore, 16 genomes had duplicate spacers within their genome with a total of 50 duplicate spacers across all the GBS genomes studied.

### Phylogenetic Trees

Two separate phylogenetic trees were constructed for 16S sequences, and cas9 of the selected genomes (Figure 3). The 16S phylogenetic analysis showed all sequences clustering into 2 major clades based on their CRISPR-Cas status. This close clustering of strains may be indicative of close intra-genus relationship among them. Cas9 phylogenetic tree showed clustering of the strains into 3 major clades grouped on the variations seen in their respective genes (Supplementary figure 1).

**Figure 3:**
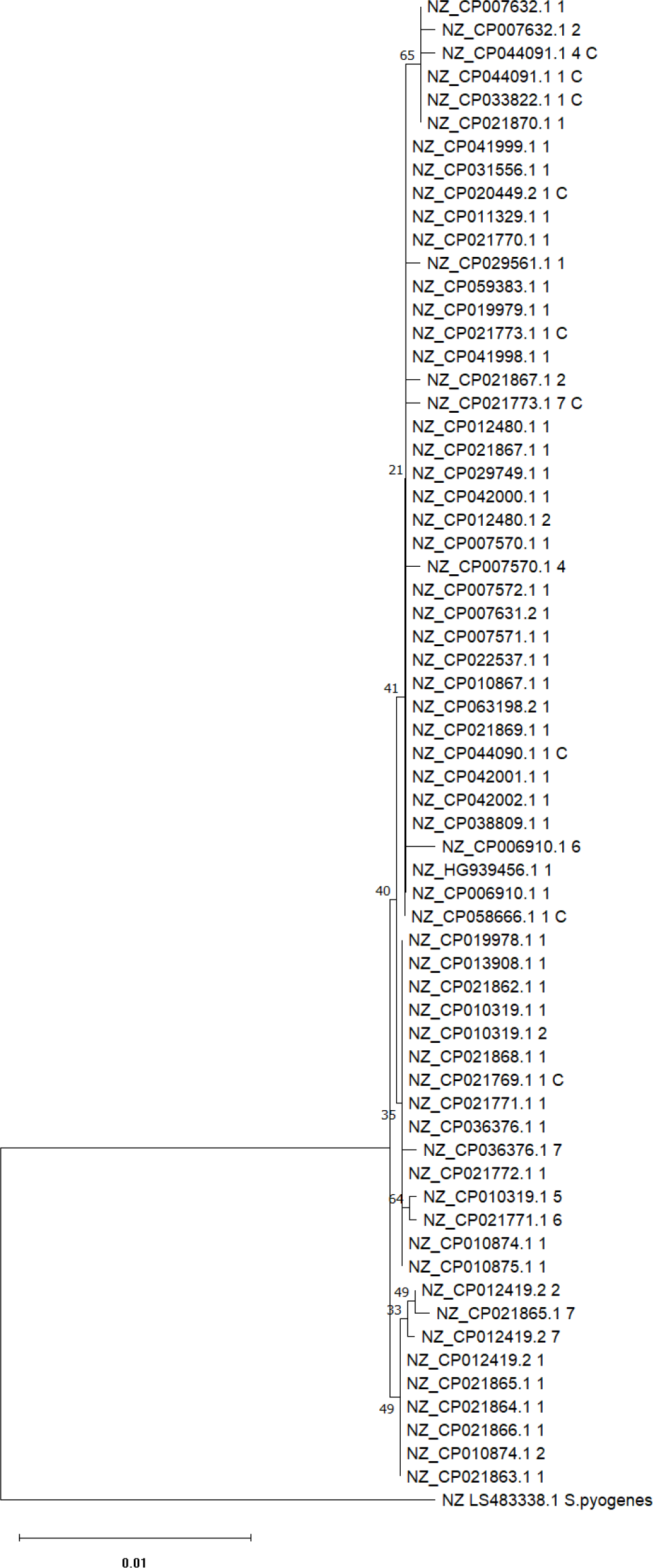
Phylogeny of GBS used in this study. The tree was based on 65 non-redundant complete 16S rRNA sequences from 51 species of GBS. *S. pyogenes* was taken as the outgroup. Numbers next to nodes indicate bootstrap values (%) based on 1000 iterations. Branch length scale indicates the number of substitutions per site. The phylogeny tree was constructed in MEGA10 using the maximum likelihood method.

## DISCUSSION

In this study, we investigated the CRISPR–Cas systems in the GBS genomes isolated from humans’ origin to gain insights into the occurrence, diversity, and features of its adaptive immune system. GBS had a high frequency of occurrence of the complete CRISPR–Cas system (91.4%). This is comparable to the reported prevalence of complete CRISPR loci for Streptococcus genera [9]. High CRISPR-Cas prevalence has been attributed to high viral abundance coupled with lower viral diversity in the ecosystem [20]. Bacterial CRISPR-Cas systems have been associated with interaction of pathogens with host cells, immune evasion and other bacterial virulence [21]. Interestingly, contradictory functions have been reported on the functioning of CRISPRs. Short or complete absence of CRISPR arrays have led to increased pathogenicity as seen in gastroenteritis causing *Campylobacter jejuni* strains [8], while cas genes has been shown to enhance virulence in *S. agalactiae* mutant studies [22]. On the one hand, CRISPR-Cas system may lessen the potential virulence by preventing MGE from introducing new virulence genes, while on the other hand, CRISPR-Cas may enhance virulence by regulating gene expression and promoting host colonization. GBS expresses various surface and secreted virulence factors to colonise and infect neonates, which also supports survival in the bloodstream.

The CRISPR–Cas systems are classified into two classes, Class I and Class II, 6 types and 33 subtypes based on the crRNA–effector complex [23]. The genera *Streptococci* fundamentally harbor type I, type II and type III CRISPR-Cas systems in addition to the individual type V and unknown CRISPR loci [24]. The type II system is involved in pathogenesis, quorum sensing, invasion and stress response among others while type I systems drives DNA targeting and cleavage associated with antiviral defense. Type III systems provides transcription-dependent immunity against diverse nucleic acid invaders [25]. In our study, out of selected 52 genomes, 51 genomes contain CRISPR arrays with a total of 75 CRISPR arrays detected and 17 strains possessing multiple of these CRISPR arrays further classified into Type II, Type I or orphan CRISPR types.

A majority, 29 genomes (55.76%) of the CRISPR–Cas systems of the GBS genomes were of Type II-A, while 15 (28.84%) genomes contained both Type II-A and I-C type of the CRISPR–Cas system. This composition is similar to the that of CRISPR-Cas of other *Streptococcus* species like *S. canis* [26] and *S. pyogenes* [27]. The type I-C in GBS contains seven cas genes (cas3, cas5c, cas8c, cas7, cas4, cas1 and cas2) similar to the ones found in S. pyogenes [27]. Cas9 was found in all 51 genomes. The recent studies indicates that Type II CRISPR-associated protein 9 (cas9) influenced virulence in GBS strains [28,29]. The virulence factors of GBS have been implicated in vaginal colonization and invasive disease through Cas9 based regulators [29].None of the genomes contained any transposon or retrotransposon elements in the CRISPR loci.

Interestingly, we are also able to detect anti-CRISPRs from GBS, which contributes to CRISPR-Cas system diversity and which also prevents genome editing. Two types of Anti CRISPR (Acr) regions were detected from selected strains as AcrIIA21 being present in 26 strains whilst AcrIIA18 in just GBS1-NY strain. AcrIIA21 exhibits broad spectrum action by inhibiting *Streptococcus pyogenes* Cas9 (SpyCas9), *Staphylococcus aureus* Cas9 (SauCas9), and *Streptococcus iniae* Cas9 (SinCas9), exhibiting high efficacy against SinCas9 [30]. An in depth understanding of its mechanism remains elusive. Furthermore, the modulation of Cas9 through sgRNA has also been reported from AcrIIA17 and AcrIIA18. The AcrIIA18 performs Cas9-dependent truncation of sgRNA which lead to generation of a shortened sgRNA which are incapable of triggering Cas9 activity [31].

CRISPR repeats are known to produce hairpin loops like secondary structure owing to its palindrome repeats. The stem-loop structure of DRs are known to facilitate the interaction between spacers and cas proteins. An investigation of the RNA secondary structures and their MFE values indicated that all but one DRs could form stable structures with ΔG < −10 kcal mol^−1^. DR1, DR3 and DR5 had lower MFE values in comparison to DR5, DR6, DR7, DR8 and DR9. Studies indicate that active CRISPR arrays tend to be long due to the continuous acquisition of spacers [32]. In this study, a maximum of 31 spacers were present in CRISPR loci indicating an active system. The average spacer length in the GBS genomes was 39 bp. In comparison, some genomes like that of *E. coli* contains an average length of 31 bp while it was found to be between 28 and 32 nucleotide bp length in *S. thermophilus* [33]. Studies indicate that CRISPR systems containing spacers of length >30 bp are more active than loci with shorter spacer lengths and more spacers allow bacteria to mount a better defense against viruses [34]. Many of the geographically close strains carried a CRISPR cassette with diverse spacers. Such observations have recorded earlier from *S. thermophilus* where spacer hypervariability has been directly linked to phage exposure [35]. Some of the spacers within the CRISPR loci were duplicated within the genome, the exact significance of this is not clear. Further experimental evidences are needed to investigate the functioning of the CRISPR–Cas systems on gene expression and regulation especially during host-pathogen interaction in GBS genomes.

## CONFLICT OF INTEREST

The authors declare that there is no conflict of interest.

## ACKNOWLEDGMENT

R. P. Shastry was supported by DST-SYST, Government of India, New Delhi (SP/YO/2019/1046).

## DATA AVAILABILITY STATEMENT

The sequence data used in this work were obtained from NCBI. The authors declare that all data supporting the findings of this study are available within the article and its supporting Information files.

## AUTHOR CONTRIBUTIONS

SDG, RPS, and RSPR planned the work. RSPR, RPS and SDG performed the work and wrote the manuscript. DP helped in data curation. PKS gave critical comments and helped in the editing. All authors contributed intellectually, and edited/reviewed the manuscript. All authors have read and agreed to the published version of the manuscript.

## SUPPLEMENTAL INFORMATION

Supplemental information for this article is available online.

## Notes

### Competing Interest Statement

The authors have declared no competing interest.

